# Neural activity in human eyeblink conditioning: an optically pumped magnetometer-based MEG study

**DOI:** 10.1101/2022.10.19.510727

**Authors:** Chin-Hsuan Sophie Lin, Tim M Tierney, Stephanie Mellor, George C O’Neill, Sven Bestmann, Gareth R Barnes, R Chris Miall

## Abstract

There is a profound lack of electrophysiological data from the cerebellum in humans, as compared to animals, because the pervading perception is that it is difficult to record human cerebellar activity non-invasively using magnetoencephalography (MEG) or electroencephalography (EEG). We used on-scalp MEG based on optically pumped magnetometers (OP-MEG) to detect learning related cerebellar signals in a classical eyeblink conditioning paradigm. In four healthy human adults, we observed unconditioned stimulus-related evoked responses locating in the cerebellum. These evoked responses diminished during the acquisition of conditioned responses, which corresponds with previous observed changes of Purkinje cell activities in animals. We also observed evoked responses immediately before conditioned blinks in 3 out of 4 participants, which were also located in the cerebellum. We discuss the potential nature of these cerebellar evoked responses and future applications of OP-MEG to the studies of human cerebellar physiology.

## Introduction

An accumulating body of literature from the past three decades has provided strong evidence that the human cerebellum supports a variety of tasks, from sensorimotor learning (Sokolov et al., 2017), event timing (Lusk et al., 2016), language processing (Mariën & Borgatti, 2018; Schmahmann & Sherman, 1998) to social cognition (Van Overwalle et al., 2020). Advances were made possible through – among other methods - detailed structural magnetic resonance imaging (MRI) and symptom mapping in subjects with focal cerebellar lesions (Bodranghien et al., 2016; Timmann et al., 2009). Functional magnetic resonance imaging (MRI) in healthy subjects has also provided key insights of cerebellar-cortical pathways that mediate a wide range of motor and non-motor behaviours (Buckner, 2013). Computational models have been informative for delineating the cerebellar learning algorithms and for generating testable hypotheses for empirical investigation (Raymond & Medina, 2018). However, the physiological mechanisms of these diverse cerebellar functions remain topics of ongoing debate. It is especially noticeable that there is a profound deficiency of electrophysiological knowledge of the cerebellum in humans, as compared to animals, because the pervading perception is that it is difficult to non-invasively record human cerebellar activity using MEG or EEG.

One model case that demonstrates the knowledge gap between animal and human cerebellar physiology is eyeblink conditioning. In the classic delayed eyeblink conditioning procedure, a neutral conditioned stimulus (CS), such as a tone or a light, is paired with a blink-eliciting air-puff unconditioned stimulus (US) with the CS-US interval being fixed and the two stimuli co-terminating. Over time, the temporal association between the CS and US is acquired and learning is manifested by a conditioned response (CR) - blinks which start during the CS and generally peak at the arrival of US. It is now well known that in both animals and humans, associative learning happens within the cerebellum (review for animal see Freeman, 2015; for human see Gerwig et al., 2007). Experimental lesions either in cerebellar lobule VI or the interposed nucleus of animals can impair the acquisition and retention of conditioning. Human patients with cerebellar pathologies are also deficient in the capacity to acquire eyeblink conditioning. Detailed lesion-symptom mapping (Timmann et al., 2009) and functional MRI studies (Cheng et al., 2008; Dimitrova et al., 2002; Thurling et al., 2015) identified lobule VI and the interposed nuclei being the eyeblink conditioning regions in humans, indicating the same functional anatomy in animals and humans.

However, our understanding of the neural mechanisms of eyeblink conditioning came almost exclusively from animal recordings. A rich body of unit and local field recordings showed that the US is transmitted to Purkinje cells via climbing fibres that originate in the inferior olive. The CS is transmitted mainly via mossy fibres that synapse on granule cells. The axons of granule cells become parallel fibres which synapse on Purkinje cells. Thus, CS and US signals converge on Purkinje cells, the sole output of the cerebellar cortex. Mossy and climbing fibres also have collaterals to deep nuclear neurons, which are the only downstream neurons of Purkinje cells. It is generally agreed that the interplay between Purkinje cells and nuclear neurons is at the heart of learning. Purkinje cells are intrinsically active, firing simple spikes at a fast and regular rate (30-150 Hz generally, up to 250 Hz)(Armstrong & Rawson, 1979; Kahlon & Lisberger, 2000), and are further driven by prolific numbers of parallel fibres. However, each single climbing fibre has more than 1000 synapses on a Purkinje cell, forming a powerful excitatory connection. Therefore, in naïve animals, cerebellar responses to US are characterised by complex spikes resulting from climbing fibre input (Jirenhed et al., 2007; Ohmae & Medina, 2015; Rasmussen et al., 2014) (called US-complex spike in this manuscript). Although powerful, these complex spikes are irregular, occurring only on a proportion of trials, with close but not uniform latencies after the US. After learning, complex spikes can also be observed early in the CS-US interval (Ohmae & Medina, 2015) (called CS-complex spike in this manuscript). Importantly, the occurrence of CS-complex spikes is accompanied by the brief suppression of simple spikes and the pause and subsequent excitation of nuclear neurons(ten Brinke et al., 2015). Nuclear output neurons then activate downstream motor nuclei driving eyelid closure and result in blinks as a CR. In addition, nucleo-olivary connections inhibit the inferior olive and thus US-complex spikes (Nicholson & Freeman, 2003). There are still fierce debates about the exact roles and importance of each anatomical site and their corresponding activities in the acquisition and maintenance of conditioning. For example, the mechanism that triggers simple spike depression and interaction between simple spikes and complex spikes during learning are both topics of active research (Rasmussen, 2020). The importance of CS-complex spike and simple spike depression to CR generation are also under scrutiny (ten Brinke et al., 2019). Electrophysiological studies nonetheless provide rich knowledge about eyeblink conditioning-related plasticity at circuitry and cellular levels.

In comparison to animal studies, even though clinical studies have strongly supported the key role of the cerebellum, direct evidence of neurophysiological changes to conditioning in humans is largely lacking. Identifying electrophysiological markers in humans is necessary for several reasons. First, while fMRI aims to provide a proxy of neural activity by measuring metabolic changes, the nature of haemodynamic responses means it is technologically infeasible to investigate the precise temporal properties of cerebellar activity using fMRI. There is also increasing evidence showing that fMRI is a better proxy of synaptic responses to inputs than spiking activity output in the case of the cerebellum (Lauritzen et al., 2012; Mathiesen et al., 1998). Therefore, identifying learning related neural activity is likely to be a blind spot of cerebellar fMRI. This means we need direct measurement of spiking activity to compare animal and human studies. Second, eyeblink conditioning undergoes changes across the lifespan (Solomon et al., 1989; Woodruff-Pak & Thompson, 1988) and in various neurodegenerative (van Gaalen et al., 2019; Woodruff-Pak, 2001) and neurodevelopmental disorders (Reeb-Sutherland & Fox, 2015). Thus, electrophysiological markers of eyeblink conditioning may well have wider clinical implications. Importantly, there is an increasing awareness that impaired eyeblink conditioning often starts in the pre-clinical stage of neurodegenerative disorders (van Gaalen et al., 2019; Woodruff-Pak, 2001). Therefore, electrophysiological markers can enhance our chances of early detection and intervention of these diseases.

Two recent studies discussed the feasibility of recording cerebellar electrical signals extracranially. Anderson and colleagues (2020) revisited a small number of reports which have detected cerebellar activity in M/EEG while Samuelsson and colleagues (2020) simulated M/EEG signals using a realistic cerebellar model based on high resolution MRI. Both studies concluded that cerebellar activity should be detectable in M/EEG. Indeed, the new type of on-scalp MEG system based on optically pumped magnetometers – OP-MEG (Boto et al., 2018) has shown great potential, enhancing the signal intensity and spatial resolution of deep brain areas including the cerebellum. And a recent EEG study has identified cerebellar activity in a classical conditioning paradigm using the otolith-evoked blink reflex (Todd et al., 2021). Given the complementary distribution of sensitivity for MEG and for EEG over the cerebellar cortex (Samuelsson et al., 2020), we believe there is scope for additional MEG focused work in this area.

Previously, we recorded event-related OP-MEG responses to air-puff stimulation of the eyes, in the absence of classical conditioning. We used on-scalp OPM sensors placed over the posterior cranium (Lin et al., 2019), and source analysis indicated the responses most likely originated from the cerebellum. In this study, we paired air-puffs (US) with auditory tones (CS) and record changes of OP-MEG in accordance with conditioning. We then infer the nature of these OP-MEG responses based on the changes we expect from animal recordings. First, we expected to see reduced US-evoked responses, irrespective of the degree of eyelid closure at the time of US. Second, in the CS-US interval, we expect change due to CS-related complex spikes concurrent with simple spike depression in the cerebellar cortex and pause and excitation in the nucleus neurons. Finally, there may be newly formed CS-related responses, although it is unlikely that the temporospatial profile of CS-related responses will be similar to US-evoked responses.

## Methods

### Participants

Four healthy adult human participants (1 female, 3 male) aged 29-52, took part in the experiment. None of them had neurological or psychiatric disorders except for participant 4 who reported a history of common migraine, which was not active and not treated with any pharmaceutics during the study. Two participants (participants 1 and 2) took part in our previous study (Lin et al., 2019) so they were experienced with the unpaired air-puff and sound-tone stimulation used in the protocol; the other two participants were naïve to the stimulation conditions. Specifically, three participants were naïve to the paired CS-US stimulation that would induce eyeblink conditioning, whereas participant 1 had experienced one session of paired stimulation, about 7 months prior to these recordings.

The study conformed to the standards set by the Declaration of Helsinki, except for registration in a database. The protocol was approved by the University College London Research Ethics Committee and the University of Birmingham Research Ethics Committee. Written informed consent was obtained from all participants. The experiments took place at the University College London.

### Eyeblink conditioning paradigm

**Figure 1A** shows the experimental setup. We used a standard delayed conditioning protocol developed by Gormezano & Kehoe (1975). The US was a brief air-puff delivered through a nozzle mounted on the helmet worn by the participant. The nozzle was connected to a pressurised air cylinder (1 Bar) through a 10 m semi-rigid plastic tube (2mm internal diameter). Under the control of a bespoke pneumatic valve controller (Leonardelli et al., 2015), the arrival time of the air-puff was estimated to lag the valve opening by 33 ms, calibrated off-line using a microphone, and was relatively insensitive to the nozzle-to-eye distance over a limited range (∼1 ms/cm). The nozzle directed the air-puff to the outer canthus of the left eye at approximately 2-4 cm from the eye. The distance was individually set such that the puff was able to evoke a visible blink after each delivery, but without causing discomfort. The CS was a 550-ms binaural tone (2800 Hz). There were two phases to each experiment, a baseline and an acquisition phase. Specifically, participants received a baseline block of trials followed by four acquisition blocks, each block lasting approximately 12 minutes; the interval between blocks was ∼1 minute. The baseline block constituted of 200 trials: 140 US-only trials, 50 CS-only trials, and 10 CS-US paired trials, with the air-puff triggered 500 ms after tone onset. The acquisition block also constituted of 200 trials: 190 CS-US trials and 10 CS-only trials. In every block, trial order was randomized in sets of 20. Every trial began with a random wait of 1-2.5 seconds to avoid habituation and anticipation; inter-stimulus intervals averaged to 3.6 s.

**Figure 1.**
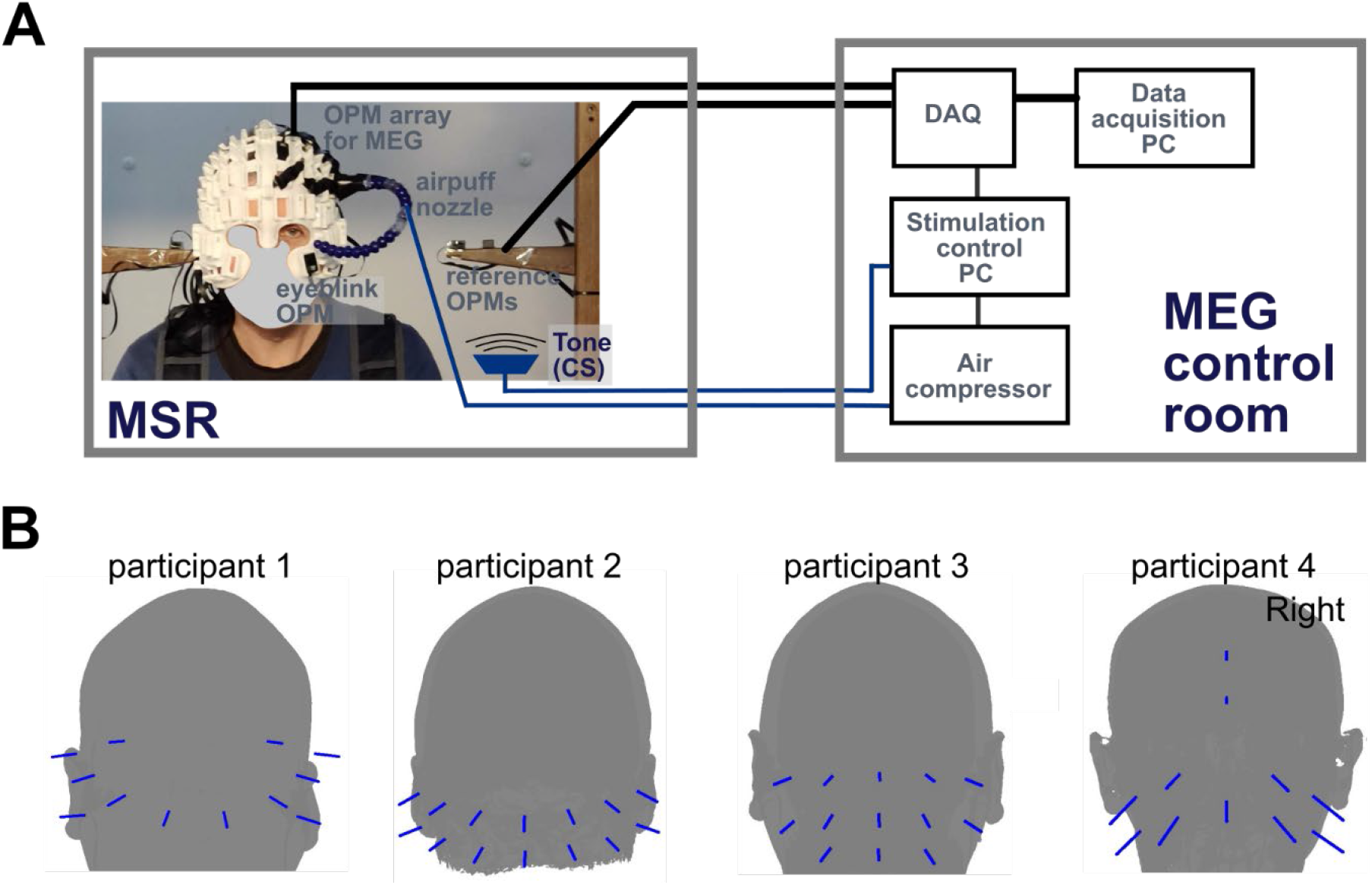
**A** Eyeblink conditioning experimental setup. The participant was seated inside a magnetically shielded room (MSR) wearing a customized ‘scannercast’. OPM sensors were inserted into slots covering the cerebellum. One more sensor was placed in the left infra-orbital slot to measure eyeblinks. Air-puffs were delivered to the outer canthus of the participant’s left eye via a nozzle connected to a bespoke air compressor. Both the air compressor and tone (CS) delivering speaker were controlled by a PC, which ran the custom-built experimental code. The data acquisition (DAQ) system recorded and synchronised OPM and trigger signals and then sent data to a data acquisition PC in real time. **B** Posterior sensor positions for the OP-MEG recordings. Positions on participant-specific ‘scannercasts’ that are most close to the cerebellum among all available slots were chosen based on each individual’s T1-weighted brain MRI. We have excluded sensors showing excessive noise in the data pre-processing stage, i.e. only OPMs used for MEG analysis are shown.

### OP-MEG data recording

During the experiment, biomagnetic signals, as well as stimulus timing signals and eyeblink responses were recorded using the OP-MEG system at the Wellcome Centre for Human Neuroimaging (University College London, UK). The use of an OP-MEG system for the eyeblink paradigm has been previously described in detail (Lin et al., 2019). Briefly, the system used 15-20 QuSpin optically-pumped magnetometers (OPMs) (QuSpin Inc., Louisville, CO, USA) (Shah, Osborne, Orton, & Alem, 2018). Each OPM sensor served as a one MEG channel recording electromagnetic signals radial to the scalp. Sensors were mounted on individually tailored rigid sensor helmets (Meyer et al., 2017), with sockets around the outer surface to hold the OPM sensors. The inner surface of each ‘scannercast’ helmet was based on each participant’s scalp extracted from a T1-weighted MRI image. Therefore, sensor sockets are in the same coordinate space as the MRI image of the brain. This means sensor positions and orientations were accurately co-registered to individual participant’s brain anatomy. Here, with limited numbers of sensors available, we used the MRI-derived 3D mesh of each individual’s brain surface and of the helmet to select sockets positioned proximal to the cerebellar cortex. Across the 4 participants, 12 to 14 posterior sensors were used. Additionally, one sensor was placed in the left infra-orbital socket to detect eye-blink. Due to excessive noise, 2 or 3 posterior sensors were removed from further data analysis for participant 1 and 4. This left 12, 14, 13 and 11 posterior sensors for participants 1 to 4, respectively. **Figure 1B** shows sensor position and orientation on the heads. Another four OPMs were placed close to the head to serve as a reference array providing ambient field strength and gradient for sensor calibration and interference rejection.

Recording was performed in an OPM-dedicated 4-layer magnetically shielded room (MSR) (Magnetic Shields, Ltd.; internal dimensions 3 m × 4 m × 2.2 m). Participants were seated with their head approximately central within the room during recording. The static field within the central 1 cubic metre of the OPM room is approximately 2 nT and the field gradient is approximately 1 nT·m^-1^ (Mellor *et al*., 2021). No additional active shielding was used.

OPM data were recorded using custom acquisition software, built in Labview. Each OPM channel was measured as a voltage (± 5 V, 500 Hz antialiasing hardware filter) and sampled at 6 kHz by a National Instruments analogue-to-digital converter (NI-9205, 16-bit, ± 10 V input range) using QuSpin’s adapter (https://quspin.com/products-qzfm/ni-9205-data-acquisition-unit/). This signal was then scaled by a calibration factor to represent the recorded magnetic field. The time-to-live signals triggering the air-puff valve and the audio buzzer were acquired simultaneously through the same data acquisition system and sent to the OP-MEG data acquisition PCs to align air-puff triggers and OP-MEG data.

### Data analysis

All of the data analysis was performed using SPM12 within the MATLAB environment (Release 2019b, Mathworks Inc., Natick, MA).

#### OP-MEG data pre-processing

OP-MEG data were first filtered between 5 and 80 Hz using a 4^th^ order Butterworth filter and then notch filtered at 50 Hz using a 3^rd^ order Butterworth filter to remove the power line noise. Reference noise cancellation was used to linearly regress the signal recorded by the reference array from the signal recorded at the scalp array (Boto et al., 2017; Fife et al., 1999). After reference noise cancellation, we examined the power spectrum density (psd) to identify any remaining residual noise peaks. One additional notch filter at 100 Hz was used to the acquisition phase data of one participant (participant 3) based on the psd. Epochs of data for each trial were extracted between −1000 and +1000 ms relative to air-puff onset for the US trials, and relative to the tone onset for the CS trials. Thereafter, acquisition phase data were concatenated across the four blocks. Trials from the baseline and acquisition phase were ranked separately according to signal variance, for each participant. These rankings were then used in a median absolute deviation method (Leys et al., 2013) for the rejection of outliers.

#### Eyeblink data analysis

Eyeblink responses were analysed on a trial-by-trial basis using the following custom pipeline. We began with the processed and epoched data from the infraorbital sensor, as described in the *OP-MEG data pre-processing* section. The data was then downsampled to 1200 Hz using SPM12 and low pass filtered at 100 Hz. Thereafter, we performed full wave rectification and low pass filtering at 10 Hz. Blink data outliners were then rejected using the same median absolute deviation criterion that was applied to OP-MEG data. For both baseline and acquisition phases, the magnitude of the unconditioned eyeblink response (UR) was defined as the peak after US onset. Conditioned eyeblink responses (CR) were identified in the acquisition phase using the following amplitude criteria: response onset was automatically detected as being when signal amplitude within the CS–US interval first deviated 3 SDs from the baseline level in the 500 ms before the CS (Thurling et al., 2015). In trials where pre-CS baseline was not stable and hence the no CR was identified by the onset criterion, a CR peak was identified when a response one-twentieth (or greater) of the mean UR magnitude occurred 150-500 ms after CS onset; responses <150 ms after CS onset are excluded as reflexive responses to the tone (Woodruff-Pak et al., 1996). For a CR that were identified using the peak amplitude criterion, the onset was then identified as the closest trough that preceded the CR peak. All trials were then visually inspected and implausible CRs were excluded. Trials were classified for further analysis as CR+/UR-, CR-/UR+ or CR+/UR+. Trials with no identifiable CR or UR due to excessive noise were marked as bad trials.

We calculated the percentage incidence of UR during baseline US-only trials and acquisition CS-US trials as well as the percentage incidence CR during CS-US and CS-only trials of both phases. We then compared the incidence of UR and CR between the baseline and acquisition phases to examine the effect of learning using a Wilcoxon rank sum test. CR incidence should ideally rise from 0 to 1 as conditioned responses become established while UR incidence should decrease. Friedman tests were used to analyse changes across consecutive blocks. *P* < 0.05 was considered significant.

#### Evoked response analysis

All the trials surviving blink and OP-MEG outlier rejection were used. For some analyses, CR+ and CR-trials were separated and compared. To identify evoked responses, trials were averaged, and baseline corrected to the mean of the window 100 ms prior to stimulus onset. In some analyses, to examine the effect of blinks on evoked response and to identify responses that were potentially time-locked to blinks, the data were re-aligned to blink onset, baseline corrected and averaged. Paired t-tests were used to examine if the peak amplitudes of evoked responses at each sensor were different from the 100 ms pre-stimulus/pre-CR time windows; *p*<0.05 was considered significant. The global field power (GFP) was estimated for each participant from the average signal of each experimental condition (Lehmann & Skrandies, 1980). Because field strengths (**Figure 3A & 3B**) and sensor layouts (**Figure 1B**) differed significantly across participants, to quantitatively compare field strengths of different conditions, we normalised GFP such that the maximum for each participant was 1. The GFP difference between baseline and acquisition or between CR+ and CR-trials was determined with paired t-tests at each time point with a criterion set at p < 0.05. To minimise false-positive results, we only accepted periods significantly different at >60 contiguous samples, i.e. >10 ms. For source level analysis the subject-specific latencies of identifiable evoked responses were calculated using peaks in the GFP amplitudes. Data from all posterior sensors were then averaged across a time window ± 2ms from the peak and were then used for source localisation.

**Figure 2.**
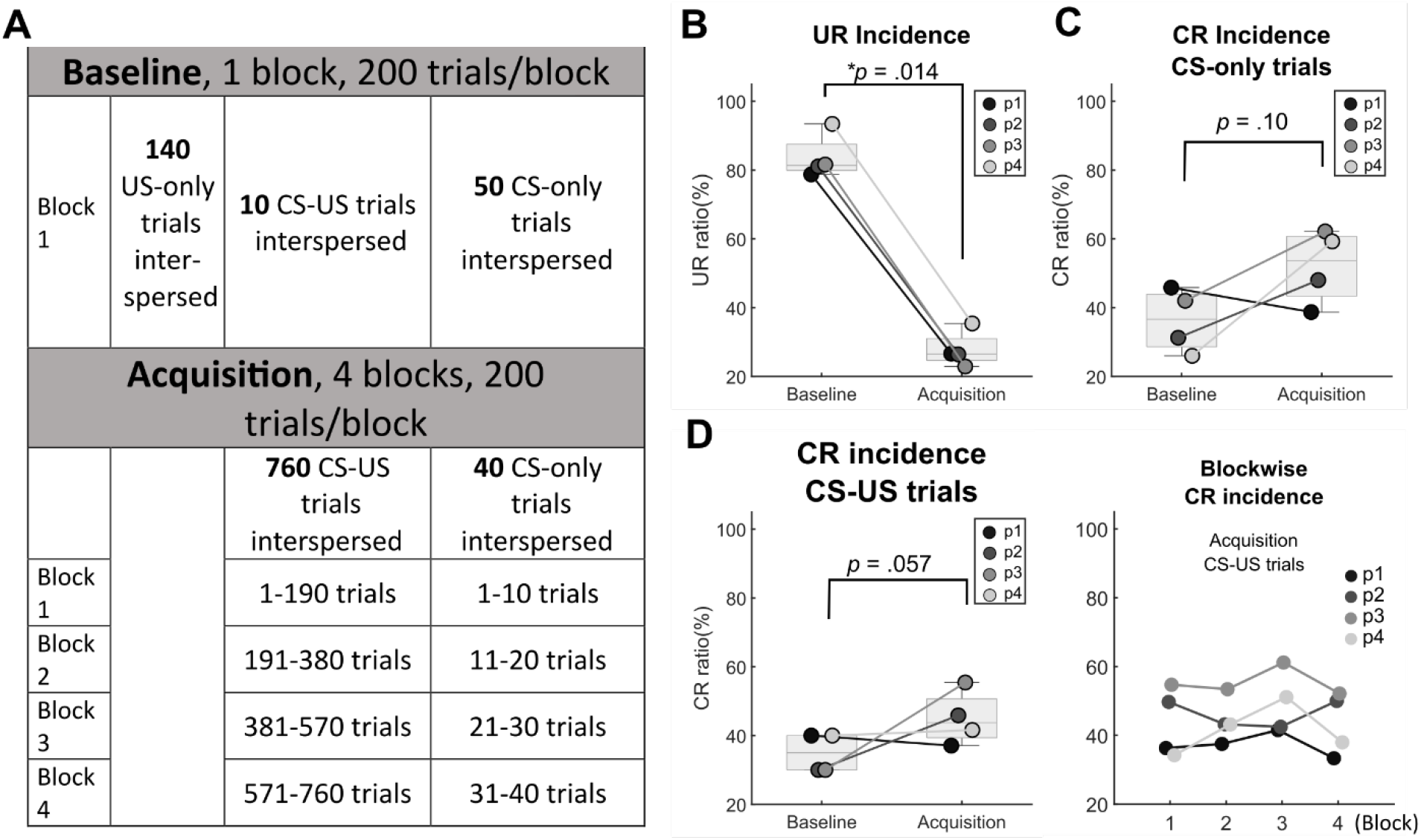
Eyeblink conditioning paradigm and blink data **A**. task structure, 1 block of baseline phase followed by 4 blocks of acquisition phase **B**. Comparing the UR incidence between baseline US trials and acquisition CS-US trials, there was a significant decrease in UR incidence from the baseline to the acquisition phase (median 81% to 27%, one-sided Wilcoxon rank sum test *p* = .014). For all the boxplots in Fig 2, the line inside the box is the median of each condition. The lower and upper quartiles are shown as the bottom and top edges of the box, respectively. Individual data points are displayed in participant specific grey dots. “p1” = participant 1 and so on. **C**. In 3 out of 4 participants, CR incidence of CS-only trials was higher in the last block of the acquisition phase, compared to the baseline. Median CR incidence of CS-only trials increased from 37% during the baseline to 54% in the last block of the acquisition phase although statistically not significant (one-sided Wilcoxon rank sum test *p* = .10) **D**. For the CS-US trials, median CR incidence increased from 35% in the baseline to 44% in the acquisition phase with a marginal significance (one-sided rank sum test, *p* = .057, left panel). Within the acquisition phase, there was no significant difference across blocks (Friedman test *χ*^*2*^(3) = 2.1, *p* = .55, right panel).

**Figure 3.**
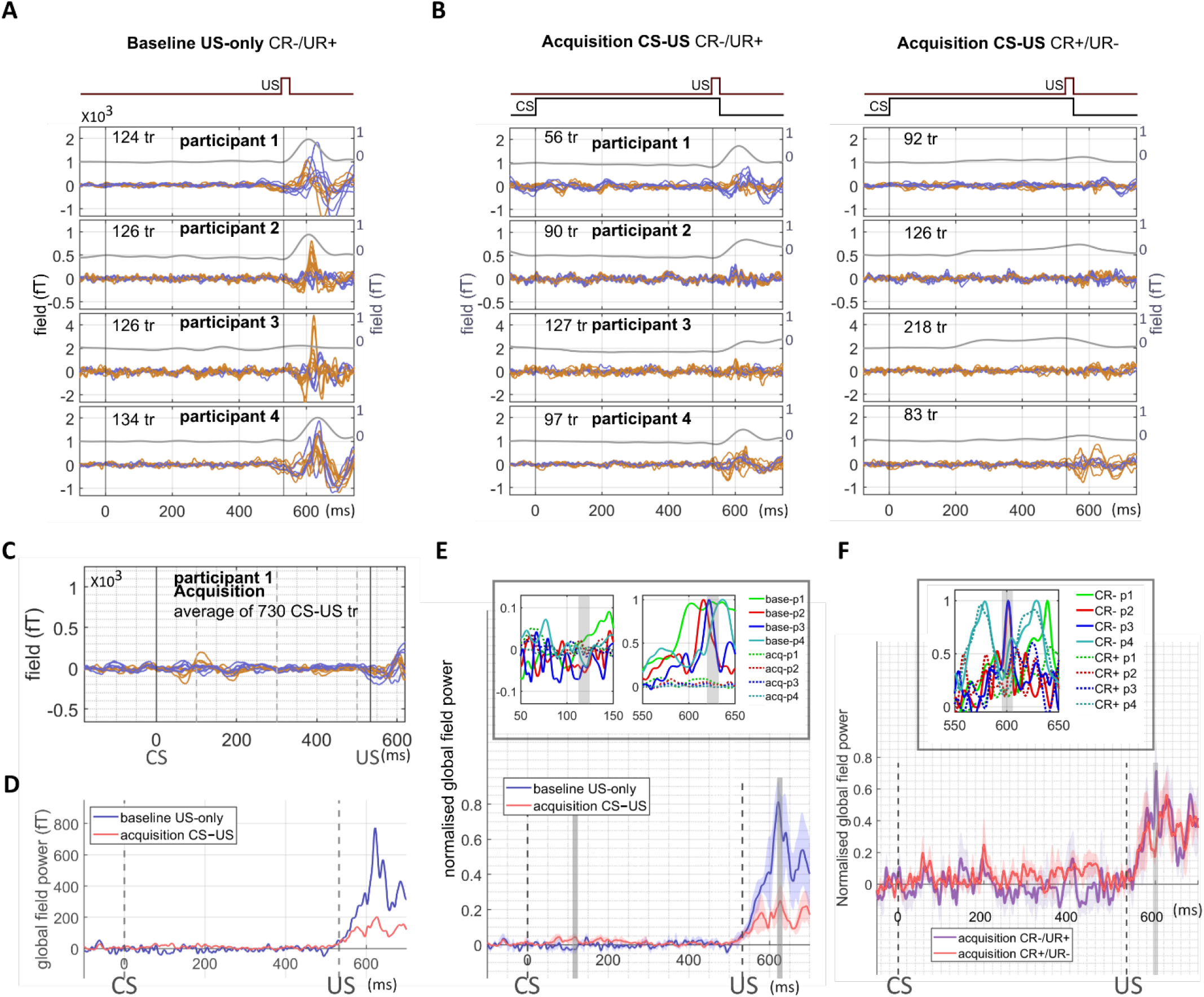
Average sensor level response and global field power. Sensor-level mean blink and evoked responses in baseline **A**. and acquisition phases **B**. Each trace corresponds to the average signal for one sensor, including the blink response from the infraorbital sensor (grey line) and evoked responses of sensors over the posterior cranium (blue and orange lines represent left and right sensors respectively) of 4 individuals. In the acquisition phase, trials were averaged separately based on whether there was an unconditioned-only blink (CR-/UR+; **Fig 3B**, left panel) or conditioned-only blink (CR+/UR-; **Fig 3B**, right panel). Note that the scales of the y-axis differ between participants but are the same between the baseline and acquisition phase in order to best present learning related OP-MEG changes. **C**. The OP-MEG of all CS-US trials aggregated and averaged. In participant 1, two dipolar responses were seen 50-150 ms post-CS. i.e. a time window similar to baseline evoked responses. No visible dipolar response seen in the other three participants (figures not shown). Note that the y-axis scale is different from **Fig 3A & B** for better data visualisation. Blue and orange lines represent left and right sensors respectively. **D**. Group means of absolute global field power (GFP) for baseline (dark blue line) and acquisition (red line) conditions. **E**. Group means of normalised GFPs for baseline (dark blue line) and acquisition (red line) conditions with shaded standard error regions. Statistical significances between the two conditions are highlighted (grey vertical bars; p < 0.05 for regions exceeding 60 contiguous samples i.e. >10 ms). At 113 ms after the onset of CS, a higher level of activation in the acquisition phase was found. At 620 ms after the onset of CS (87 ms after the onset of US), a lower level of activation was found in the acquisition phase data. The top inset plot zooms into these two time windows, showing individual GFPs, with solid lines representing baseline data and dot lines representing acquisition data. Note that only US trials were included in the baseline average. **F**. Group mean GFP comparison between CR-/UR+ and CR+/UR-trials in the acquisition phase. At 595 ms after the onset of CS (= 62 ms after the onset of US), a lower level of activation was found in CR+/UR-trials (red line), as compared to CR-/UR+ data (violet line). The top inset plot zooms into the significant time window, showing individual GFPs, with solid lines representing GFPs of CR-/UR+ trials and dot lines representing CR+/UR-trials.

#### Induced responses analysis

For induced spectral power changes, single trial time-frequency (TF) decompositions in sensor space were calculated for each subject using a Morlet wavelet transform (Tallon-Baudry et al.,1998) and then averaged across trials and the power of the evoked responses subtracted out. We used CR trials in the acquisition phase for this response analysis. Data was down-sampled to 1200 Hz before wavelet transform. The wavelet transform was calculated for each time-point between −1000 and +1000 ms, with 76 scale bins corresponding to frequencies between 5 and 80 Hz. For each trial and frequency, the mean power in the interval 100 ms before stimulus onset was considered as a baseline. The power change in each frequency band post-stimulus was expressed as the relative percentage change from the baseline. Paired t-tests were used to analyse the changes of power and p<0.001 was considered significant.

#### OP-MEG source localisation - dipole fitting

We evaluated the locus of the average evoked response using a dipole fit analysis. The volume conductor model was the Nolte single shell model (Nolte, 2003), implemented in SPM12, using the inner skull boundary from the individual T1-weighted MRI.

We reconstructed sources of the evoked field data for each subject using the SPM implementation of equivalent current dipole fitting (Kiebel, Daunizeau, et al., 2008). In brief, the inversion scheme assigned initial means and variances of dipole positions and moments (also called ‘priors’). The final dipole locations and moments were optimised using variational free energy (Friston et al., 2007; López et al., 2014; Penny, 2012). Free energy quantifies a model’s ability to explain the data by maximising the likelihood while penalising model complexity. Importantly, models with different anatomical priors can be compared using free energy to select the best dipole fit.

We specified the initial mean locations of eight single dipole models based on the literature, which included the face area of the right (model 1) and left (model 2) somatosensory cortex S1 (Nevalainen, Ramstad, Isotalo, Haapanen, & Lauronen, 2006); lobule VI in the right (3) and left (4) cerebellar cortex; (Cheng et al., 2008); right (5) and left (6) medial parieto-occipital areas, which have been shown to present blink-related activity (Bardouille et al., 2006; Liu et al., 2017). We also included the right (model 7) and the left (model 8) interposed nuclei of the cerebellum (Thurling et al., 2015). The standard deviation of each prior dipole location was set to 10 mm; the mean and standard deviation of each prior moment were assigned as 0 and 10 nA·m, respectively. To avoid local maxima, 50 iterations, with starting locations and orientations (i.e. the means and standard deviations of prior locations and moments) randomly sampled from prior distributions, were carried out for each model (50 × 8 = 400 iterations for each subject). We selected the model parameters corresponding to the highest free energy value across all iterations; the maximal free energy values were then averaged across subjects.

#### OP-MEG source localisation - beamforming

We used the scalar version of a linear constrained minimum variance beamformer algorithm implemented in the DAISS toolbox for SPM (https://github.com/spm/DAiSS) to localise the source of induced spectral power changes. The volume conductor model was the Nolte single shell model (Nolte, 2003), implemented in SPM12, using the scalp boundary from the individual T1-weighted MRI. We used an individualised covariance window (15 to 35 Hz for participants 1 and 2 and 5-35 Hz for participant 3. No induced response was observed in participant 4 so we did not conduct beamforming analysis for this participant), and we contrasted a time window of +100 to +483 ms against a window of −284 and 99 ms, relative to stimulus onset. The regularization rate λ was set to be 10 (Barnes et al., 2004). The source orientation was set in the direction of maximal power. The reconstruction grid spacing was 10 mm.

## Results

### Behavioural data

Moderate conditioning effects could be observed in the blink responses recorded from the infraorbital sensor. First, in the baseline phase we were able to detect URs with high reliability. The median incidence of URs during US-only trials in the baseline phase was 81%, significantly higher than 27% during the CS-US trials in the acquisition phase (**Figure 2B** one-sided Wilcoxon rank sum test, *p* = .014, baseline/acquisition 78/26, 81/27, 82/27 and 93/23% for each participant). Looking next at the CS-only trials, in 3 out of 4 participants, the incidence of CRs was higher in the final acquisition block compared to the CS-only trials within the baseline although statistically not significant (**Figure 2C** baseline vs acquisition 46/39, 31/48, 42/62 and 26/59% for each participant, one-sided Wilcoxon rank sum test, *p* = .10).

CRs also occurred in about 44% of CS-US trials in the acquisition phase. The median CR incidence of CS-US trials in the acquisition phase was marginally higher than observed in the small number of CS-US trials within the baseline period. (**Figure 2D left panel** baseline vs acquisition 40/37, 30/46, 30/55 and 40/42% for each participant, one-sided rank sum test, *p* = 0.057). We did not find significant change of CR incidence in these trials across the 4 acquisition blocks (**Figure 2D right panel**, Friedman test *χ*^*2*^(3) = 2.1, *p* = .55).

In summary, while the CRs observed were higher than expected in baseline, and lower than hoped for in the acquisition phase, there was evidence of associative learning in all 4 participants.

### Evoked responses

#### Baseline phase

In **Figure 3A**, we show the average waveform of US-only trials from the baseline OP-MEG data. We found two dipolar responses at time windows of 40-65 ms and 80-115 ms after the air-puff in all 4 participants which were largely similar to what we observed previously (Lin et al., 2019). The amplitudes of both early and late peaks diminished significantly when the data were re-aligned to the onset of URs (**Supplementary Figure S1A**). Furthermore, there was no correlation between OP-MEG activity and eyelid position at either earlier or later time points. (**Supplementary Figure S1B**). These results suggest that the evoked responses were stimulus- and not response-related.

#### Acquisition phase

Compared to baseline response, diminished post-US evoked fields were observed in both early and late components in the acquisition phase (i.e. at about 40-65 and 80-115ms, **Figure 3B**). The reductions can be seen in the average of both CR-/UR+ (**Figure 3B** left panel) and CR+/UR- (**Figure 3B** right panel) trials. Therefore, this effect was not likely to be only a result of decreased response to air puffs because of the pre-emptive closure of the eyelids. When taking the average OP-MEG of all CS-US trials, other than reduced post-US evoked fields, two small dipolar responses can be seen in 50-150 ms post-CS in participant 1 (**Figure 3C**). No dipolar response was observed in the CS-US interval among the other three participants (figures not shown). The post-US response reduction can be summarised by the Grand averaged GFP waveform (**Figure 3D**). Comparing normalised GFPs between the baseline and acquisition phases, there were two time windows where statistically significant difference was found using a criterion of p< 0.05 for more than 60 contiguous samples (i.e. >10 ms). At 113-124 ms after the onset of CS, a higher level of activation in the acquisition phase was found (**Figure 3E)**. At 87-99 ms after the onset of US, activation in the acquisition phase was significantly lower than the baseline phase (**Figure 3E**). We also found a significantly lower level of activation in CR+/UR-trials, as compared to CR-/UR+ trials, at about 62-72 ms post-US (**Figure 3F**); this was close to but slightly earlier than the acquisition-baseline difference (**Figure 3E**). Thus, in the acquisition phase there was a greater GFP response after the CS, and a reduced response after the US, contingent on the CRs.

#### Detailed nature of evoked responses and source analysis

The peak latencies of baseline early and late post-US components are presented in **Figure 4A** for each of the participants. Field maps (**Figure 4B**) calculated at times corresponding to the early and later peaks show quasi-dipolar patterns. Each sensor that showed a peak response statistically different from the 100 ms pre-US time window (paired-t tests) is displayed on the field map as a black dot. For both early and late peaks, significant responses were identified in more than half of the sensors in all participants (Early peaks: 9/12, 7/14, 7/13 and 10/11 sensors for each participant. Late peaks: 10/12, 10/14, 10/13 and 9/11). Free energy values for the eight single dipole models were averaged across participants, with models fitted for subject-specific 4 ms windows of early (64–68, 54–58, 61–6, and 37-41 ms; **Figure 4C** upper panel) and late responses (115– 119, 81–85, 87–91, and 101-105 ms; **Figure 4C** lower panel). The winning model for both early and late components included a cerebellar dipole. For the early peak, model with a left (ipsilateral) cerebellum lobule VI dipole was the best but not significantly better than the model with a right lobule VI dipole (ΔF=0.42, **Figure 4C** upper panel). For the late peak, the right (contralateral) cerebellum lobule VI model was the winning model, but not significantly better than the right interposed nucleus model (ΔF=1.41, **Figure 4C** lower panel). At an individual level, the best fitting dipoles for both early and late peaks were in the right cerebellum for early 3/4 and late 4/4 participants, while the 4^th^ participant had best fitting left-early and right-late cerebellar dipoles (**Figure 4D**). See **Table 1** for dipole positions and moments. These results suggest a strong cerebellar response that follows the US and precedes the UR eye blink.

**Table 1.** dipole locations, moments and the ΔF (F_best-model_ –F_2nd best-model_) of baseline evoked response data

|  | Baseline |  |  |  |  |  |  |  |  |  |  |  |  |  |  |  |
| --- | --- | --- | --- | --- | --- | --- | --- | --- | --- | --- | --- | --- | --- | --- | --- | --- |
|  | Early (40-65 ms) |  |  |  |  |  |  |  | Late (80-115 ms) |  |  |  |  |  |  |  |
|  |  |  | MNI coordinates |  |  | Dipole moments |  |  |  |  | MNI coordinates |  |  | Dipole moments |  |  |
| | $\Delta F$ | | X | Y | Z | X | Y | Z | $\Delta F$ | | X | Y | Z | X | Y | Z |
| Subj1 | -6.48 | R VIII | 29.58 | -56.40 | -54.56 | -17.80 | -33.79 | -19.00 | 4.77 | R VIII | 31.21 | -68.42 | -57.50 | -12.08 | 34.83 | 6.87 |
| Subj2 | -5.71 | R IX | 6.51 | -56.03 | -64.39 | 7.57 | 20.35 | 2.73 | 2.63 | R VIII | 19.04 | -63.47 | -59.89 | -21.86 | -21.79 | -7.65 |
| Subj3 | 3.69 | R VIII | 19.58 | -69.14 | -58.31 | 26.73 | 44.04 | 8.21 | -11.6 | R Crus II | 6.07 | -88.06 | -41.69 | -21.25 | 78.77 | 52.46 |
| Subj4 | 10.21 | L VIII | -25.30 | -36.40 | -55.18 | 41.56 | -17.37 | -2.67 | 9.83 | R Crus I | 48.53 | -77.45 | -38.10 | -5.35 | 30.93 | -49.61 |

**Figure 4.**
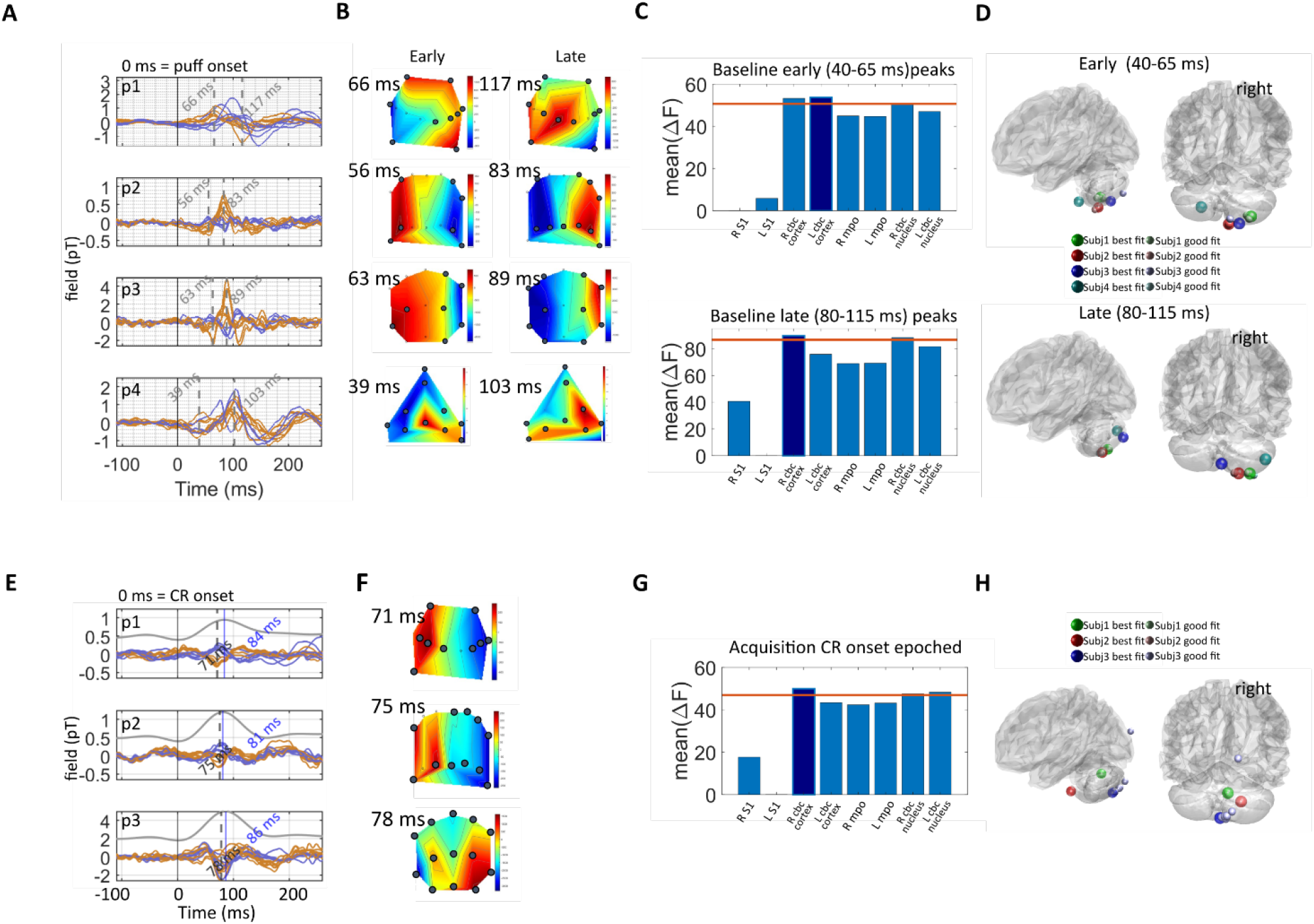
The peak latencies, field maps and dipole fit locations of evoked responses in baseline and acquisition phases. **A**. Peak latencies of the early and late post-US peaks in the baseline phase for 4 participants. Each trace corresponds to the average signal for one sensor over the posterior cranium, situated left (blue curves) and right (orange curves) of the midline respectively. “p1” = participant 1 and so on. **B**. Field maps of the evoked field at the individual-specific latencies of the two distinct peaks. On the field maps, all sensors at which the response was statistically significant compared to pre-US are displayed (black dots). Statistical significance was assessed with paired t tests comparing mean amplitudes between peak latency ±2 ms and −4 ms to 0 ms relative to air-puff contact (*p* < 0.05) **C**. Single dipole model comparison. Free energy (F) is used to approximate the model evidence of a given source solution. Bars represent the mean free energy value relative to the poorest model (which was left S1 for both early and late peaks). The left (ipsilateral) cerebellum lobule VI has the highest model evidence for the early peak and the right (contralateral) cerebellum has the highest model evidence for the late peaks. It should be noted that there were other cerebellum models for which evidence was not significantly inferior to the cerebellum lobule VI model, for both peaks. **D**. single dipole fits for each participant. Large spheres represent the source locations of fits with the highest evidence for each individual. Smaller spheres are fits which are suboptimal but not significantly so i.e. ΔF = (F_best-fit_ − F) < 3. **E**. Evoked responses of CR+ trials in the acquisition phase when trials were aligned to the onset of CRs. The same as **Figure 4A**, each colour trace corresponds to the average signal for one sensor over the posterior cranium, situated left (blue curves) and right (orange curves) of the midline respectively. Gray curve is the average blink. Note the change in scale for participant 1. Peak latencies of OP-MEG response (71, 75, and 78 ms for participant 1,2, and 3) are marked in grey texts, and latencies of blink peaks (84, 81, 86 ms for participant 1,2, and 3) in blue text. Participant 4’s data are not displayed here because there was no visible OP-MEG peak (see supporting information **Figure S4**). **F, G, H**. Field maps, single dipole model comparison and best fit locations of evoked responses identified in the acquisition phase.

Although GFP showed a small but significant increase at ∼100 ms after CS onset in the acquisition phase OP-MEG, we did not find any visible peaks in the average waveform during the CS-US interval, i.e. 0-550 ms, except in participant 1 (**Figure 3C**). To explore the possibility that some activity may be temporally correlated to CRs, as previously found in animal recordings (ten Brinke et al., 2015, 2017), we re-aligned the data using the CR onset of each CR+ trial and examined the average waveform. In 3 out of 4 participants, we identified dipolar responses peaking at 6-13 ms before blink peaks (**Figure 4E & Supplementary Figure S4**). For these 3 participants, the responses were significantly different from the pre-CR 100 ms baseline in most channels (9/12, 10/14 and 13/13 sensors; **Figure 4F**). Source inversion again identified the right cerebellar lobule VI as being the winning model (**Figure 4G**), with evidence for both right and left interposed nucleus models not significantly lesser (right VI - right interposed ΔF= 2.40; right VI - left interposed ΔF= 1.60) than for the right lobule VI. All the best model fits were found to be in the right (contra-lateral) cerebellum or brainstem (**Figure 4H & Table 2**). However, we should note that fits were also found in the occipital lobe in participant 3. See table 2 for dipole positions and moments.

**Table 2.**
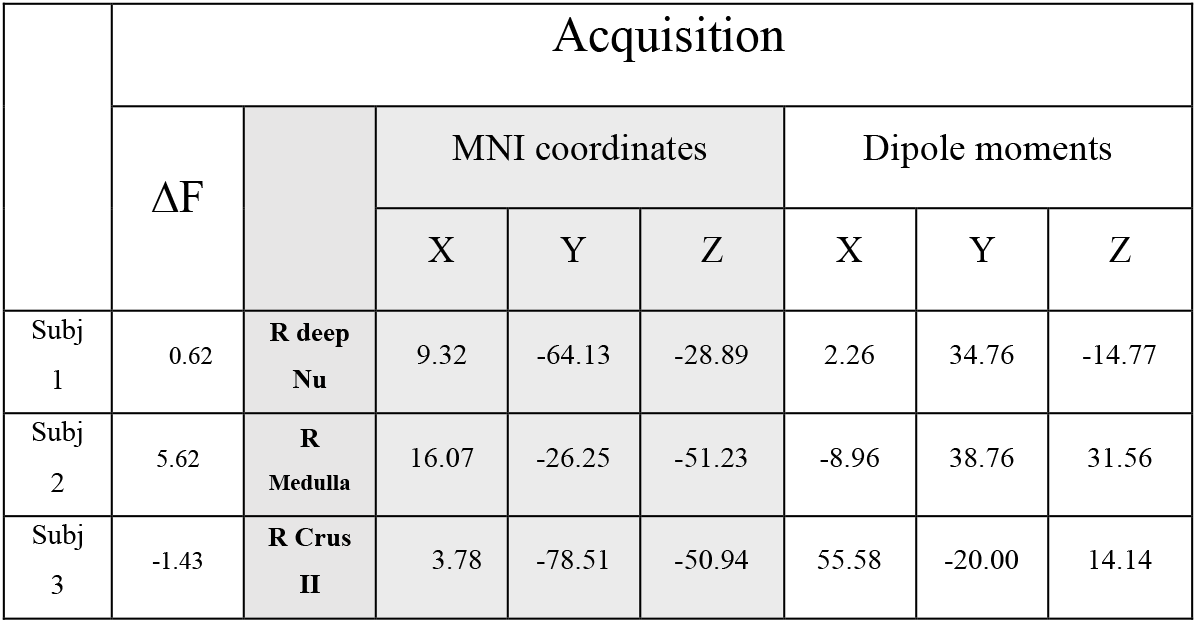
dipole locations, moments and the ΔF of acquisition phase data

This analysis reinforces the suggestion that there is an acquired post-CS response in the OP-MEG signal, that precedes the peak of the learnt CR eye blink, albeit that it appears to follow after the onset of the CR.

### Induced responses

Sensor level induced power changes in the CS-US interval of each sensor were ranked based on the mean power between 50-483 ms post-CS. The first 50 ms was excluded because neuronal modulation was not expected to occur there (Jirenhed et al., 2007). The last 50 ms (483-533 ms) of data was excluded to minimise the possibility of ranking being affected by US-related modulation. Data was then compared against the baseline using a paired t-test with a threshold of p < .0001. To minimise false activations, we only accepted responses exceeding 50 ms in duration and 5 adjacent frequencies bins (5 Hz). **Figure 5** shows the average time–frequency spectrograms (as percentage change of power from baseline) for the two sensors with the largest power change in participant 1-3. No significant power increase was found in participant 4’s data. We identified increased power in a frequency band of 5-35 Hz and a time window between 100-483 ms post-CS, with some individual differences. However, source analyses suggested the signals were from generators outside the brain (figure not shown). We consider the small number of sensors used in the experiment may have affected the performance of beamformer negatively. Noticeably, the sensors carrying the strongest signal were in the left hemisphere for 2/3 participants, unlike the right-hemisphere evoked responses for all 3 participants (**Supplementary Figure S2**). We did not find systematic suppression of sensor level responses in the time window of interest.

**Figure 5.**
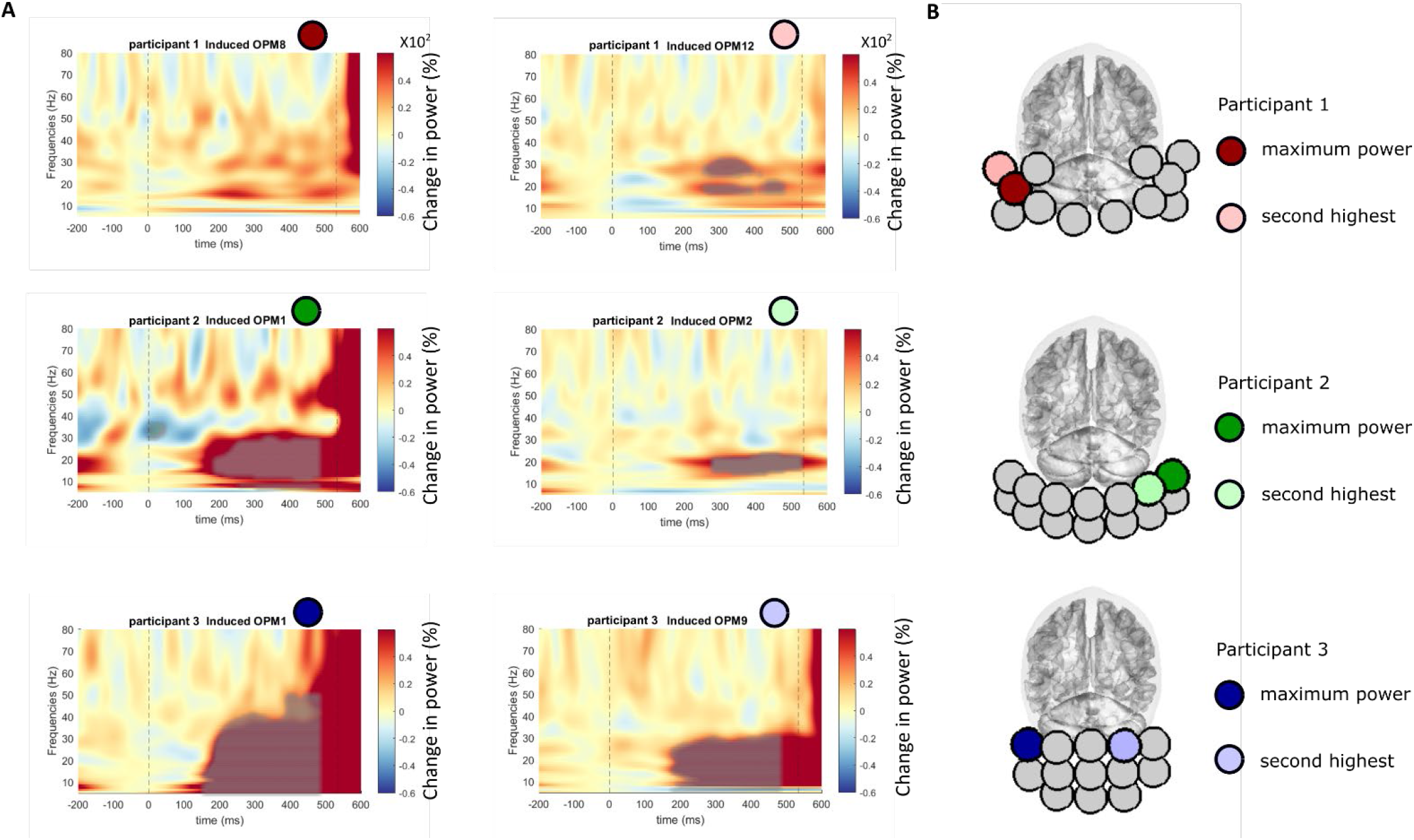
Time–frequency spectrograms showing induced power changes in the CS-US interval. **A** Each participant’s average across all trials at the two sensors with the maximal power changes are displayed. Increased activity can be observed at 15–35 Hz (5-35 Hz in participant 3), occurring from 100 ms post-CS onwards to US onset. Participant 4’ data was not presented here because no significant induced power change was found. **B**. The positions of these sensors are marked as coloured circles for each subject and mapped onto the Montreal Neurological Institute (MNI) template brain.

## Discussion

We recorded on-scalp OP-MEG in the posterior head area during the classic delayed eyeblink conditioning paradigm in 4 participants. Our aim was to identify electrophysiological markers for eyeblink conditioning in neurotypical adults. To begin with, we found multiphasic bipolar cerebellar evoked responses to the unconditioned air puff stimulus (US). We then showed that this US-related response attenuated in the acquisition phase and the attenuation was more pronounced in trials with conditioned responses (CRs). We however only found evoked responses in the CS-US interval when aligning trials to the onset of CRs. In a time-frequency analysis, there was no induced power-change response that we could attribute to a cerebellar origin.

The post-US evoked response was close to our previous findings (Lin et al., 2019), with similar latencies and a multiphasic appearance. We again localised the sources within the cerebellum. In our previous study, we showed that this response did not originate from the somatosensory cortex or the eyes. We again found these evoked responses had no temporal correlation with the blink responses (**Supplementary Figure S1**). We believe the response is most likely to be Purkinje cell activity. The dendritic trees of Purkinje cells align in the sagittal plane of the cerebellar cortex and have a high degree of synchrony within the same functional zone (De Zeeuw et al., 2011). Both features mean that Purkinje cells are likely to contribute to magnetic fields that can be detectable extracranially. In addition, the very powerful complex spike of Purkinje cells is a characteristic US-related activity (Ohmae & Medina, 2015) and so may contribute to the US-related potentials. While the US also elicits neuronal activity in the deep nuclei, post-US responses in the nuclei were expected to be enhanced rather than attenuated by learning (Nicholson & Freeman, 2000), which is at odds with the findings in this study.

Questions remain to be answered regarding the nature of some features of US-related responses. One thing that stood out is that we consistently observed multiple peaks, with early and late peaks evident in each participant. Our initial hypothesis (Lin et al., 2019) was that we would detect complex spike responses. Animal studies have repeatedly found multiphasic complex spikes – hence the name – (Eccles, 1967; Jirenhed et al., 2007; Mostofi et al., 2010; Offenhauser et al., 2005) which have functional importance to learning (Rasmussen, 2020; Rasmussen et al., 2013; Zang & Schutter, 2019). However, these bursts are typically only of 5-10 ms duration and unlikely to directly explain the longer 50-100 ms timing of our responses. Unlike complex spikes, simple spikes are typically modulated around a high baseline firing rate, and simple spike firing is known to be increased by the US (Nicholson & Freeman, 2004) immediately before US-driven complex spikes (Burroughs et al., 2017). Therefore, it is possible that the earlier component represents the modulation of simple spikes as has been suggested in a previous cerebellar MEG by (Hashimoto et al., 2003). However, after learning, the simple spike rate typically reduces prior to the CR, releasing the cerebellar nuclear cells from inhibition (Jirenhed et al., 2007; Kotani et al., 2006). Hence it is difficult to disentangle possible bidirectional modulation of the simple spike rate. An alternative explanation could even reverse these suggestions. There is some evidence of an early complex spike response followed by a later simple spike burst responsible for the termination of the eye-blink (Ohmae et al 2021). If so, one would expect the early response to decline and the late response to shift, as the CR gradually replaces the UR; we cannot confirm this with our current results.

One of the main concerns regarding our hypothesis that the post-US response is driven by complex spikes is the latencies of the peaks observed in our studies (this paper and Lin et al, 2019). In small mammal recordings, the latencies of US-triggered complex spikes are around 25 ms (Ohmae & Medina, 2015; ten Brinke et al., 2015) although they could range as far as 60 ms. Some of our responses, especially the late component, occurred much later, at 100-150 ms. Therefore, while the reduced GFP amplitudes in the acquisition phase together with a more prominent decrease of GFP in CR+ trials (**Figure 3E &F**) are consistent with the expected reduction of complex spike activity after associative learning, as evident from animal recordings (Nicholson & Freeman, 2003), our current evidence is not enough to conclude that the US-related evoked response is driven by complex spikes per se. Nor was it related to simple spike enhancement. Future studies are needed to further understand the neuronal sources of the response.

There were significant between-subject differences in the spatiotemporal distributions of these US-related responses. Comparison is challenging because the sensor layouts (**Figure 1B**) on individual scannercasts differed substantially; additionally, the number of sensors available was low, hence spatial resolution is low. We calculated the average waveform from a single sensor in each dataset, chosen to have similar on-scalp position across participants and we showed these responses were largely comparable (**Supplementary Figure S2**), with a small negative deflation followed by a larger positive and then negative peak. For future studies, we plan to build composite helmets, so that we can maintain identical sensor layout when recording different people. By doing so, we can systematically study the spatiotemporal map of the US-related waveform and build norms in the general population.

CS-related complex spikes have been reported after acquisition of eye-blink conditioning in multiple animal studies (Mostofi et al., 2010; Ohmae & Medina, 2015; ten Brinke et al., 2015), typically with a peak of complex spike probability somewhat later and broader than that following the US. However, there was no visible peak in the CS-US interval in our data, except in participant 1 (**Figure 3C**). Several reasons may be behind the result. First, learning is marked not only by CS-evoked complex spikes by also by strong simple spike suppression (Hesslow, 1994; Jirenhed et al., 2007; Rasmussen et al., 2014) together with the pause and subsequent activation of nuclear neurons (ten Brinke et al., 2015). These concurrent responses may weaken signals in extracranial recordings. Second, there was only moderate evidence of conditioned behaviour in our experiment, implying learning may have not reached an optimal level. It is suggestive that the increase in post-CS potentials (**Figure 3C**) is greatest in participant 1 who had experienced one session of paired CS-US trials several months before. It may be valuable to adopt a pre- and post-acquisition design, in future experiments, to ensure stronger acquisition. Third, many studies (e.g. Rasmussen et al., 2014) find that few Purkinje cells display enhanced complex spikes on every trial, after learning. Importantly, in their observations, simple spike suppression temporally proceeded the CS-driven complex spikes, when they were present, and was strongly correlated with the near absence of complex spikes in the later time points of CS-US intervals. Therefore, our findings raise an interesting question about how the intricate balance between enhanced and suppressed Purkinje cell activity is evident at population level, which is not easily answered by single unit recordings alone.

The nature of the small but significant cerebellar evoked responses identified in 3 out of 4 participants, when trial data were aligned to the onset of CRs, remains to be determined. The responses peaked immediately (< 10 ms) before the peak of the CR eye blink, instead of at CR onset, so they were unlikely to be CS-driven complex spikes. One potential source may be conditioning-related spikes in the nuclear neurons (Rasmussen et al., 2014; ten Brinke et al., 2017). However, the surface model we used for source reconstruction cannot differentiate between cortical and nuclear sources (**Figure 4G**). We also found neither temporal nor amplitude correlation between individuals’ overall OP-MEG and their blink data (**Supplementary Figure S3**). Correlations between neural activity and conditioned behaviours have been discovered in simple spike suppression (ten Brinke et al., 2015) and nuclear neuron facilitation (ten Brinke et al., 2017). Taking all evidence together, it is difficult to conclude on the functional implication or the neural generator of these CR-related responses.

A well-documented change of Purkinje cell activity after conditioning is the suppression of simple spikes during the CS-US interval, maximal just before the CR. The mechanism for this delayed response is still debated: it may be associated with delayed firing within the granule cell/Golgi cell/molecular layer interneuronal circuit, or to intrinsic dynamics of the Purkinje cells themselves. We analysed the induced power spectrogram in the CS-US time window. A previous EEG study observed reduction of power from beta-band (13-30 Hz), gamma-band (30-80 Hz), and up to 320 Hz that may be related to spike suppression (Todd et al., 2019). While beta and low-gamma bands are within the range of our analysis, frequencies beyond 80 Hz were not observable in the current study because of the limited bandwidth of the OPM sensors we used. Our analysis therefore searched for changes between 5-80 Hz. However, we could not find significantly decreased power in our data, and in fact 3 participants showed increase in power at 5-35 Hz and no source of theses power change was found inside the brain. Further studies are still much needed to find the neural representation of simple spike suppression after acquisition in the domain of M/EEG.

To summarise, for the first time we used OP-MEG to identify neural responses that are associated with classic eyeblink conditioning. We found attenuated US-related response after conditioning, with similarities to the complex spike firing rate changes observed in animal studies. We did not find strong evidence of other neural substrates, such as correlates of simple spike suppression or nuclear activity. Future studies could include (1) using a generic helmet to increase sample size; (2) ensuring stronger conditioning, potentially over several experimental sessions; (3) applying a high resolution cerebellar surface model that can enhance the spatial resolution of source location; (4) applying a neural mass model such as the dynamic causal modelling (Kiebel, Garrido, et al., 2008) and (5) concomitant OP-MEG/EEG recording that can help us to better understand the neural signatures for eyeblink conditioning in OP-MEG, and hence to better understand the human cerebellar computation.

## Acknowledgements

This work was supported by a Wellcome collaborative award to GRB, Matthew J. Brookes and Richard Bowtell (203257/Z/16/Z, 203257/B/16/Z). The Wellcome Centre for Human Neuroimaging is supported by core funding from the Wellcome [203147/Z/16/Z]. CSL was funded by a Wellcome grant WT212422. SM was funded through the EPSRC funded UCL centre for doctoral training (EP/L016478/1). GRB was funded through the Engineering and Physical Sciences Research Council (EPSRC) and Medical Research Council (MRC) [grant number EP/L016052/1]. RCM was funded by a Royal Society Leverhulme Senior Fellowship and by a Wellcome grant WT212422. We thank Uta Noppeney for use of the pneumatic controller to deliver air-puffs.

