## Supplementary figures for "Neural activity in human eyeblink conditioning: an optically pumped magnetometer-based MEG study"

### Supplementary Figures

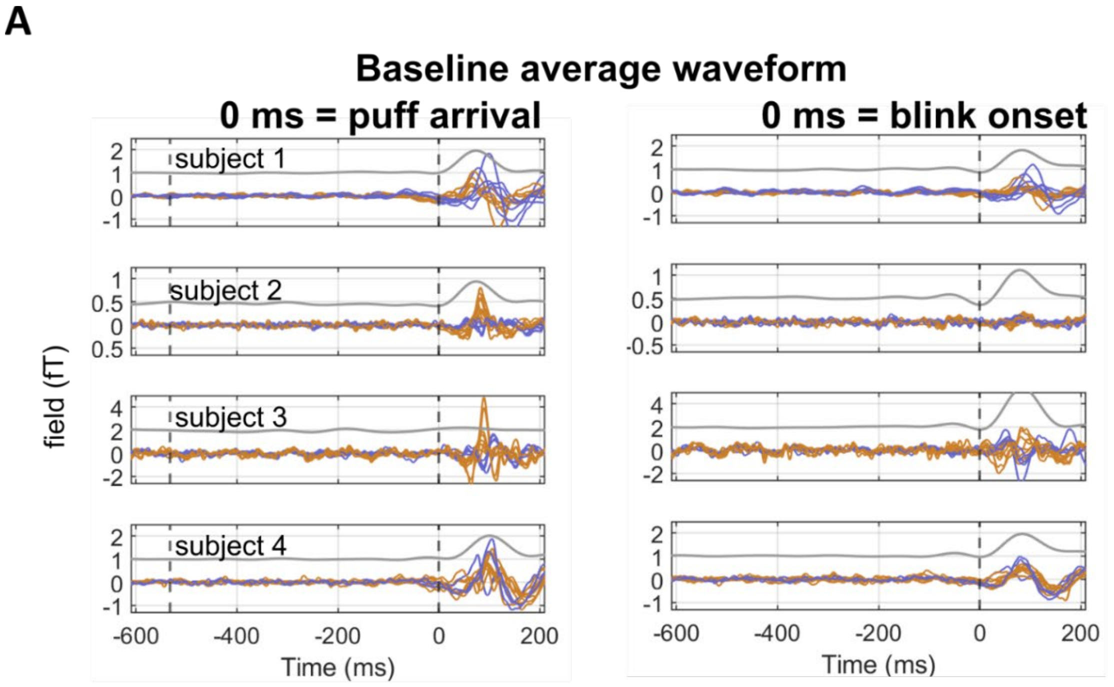


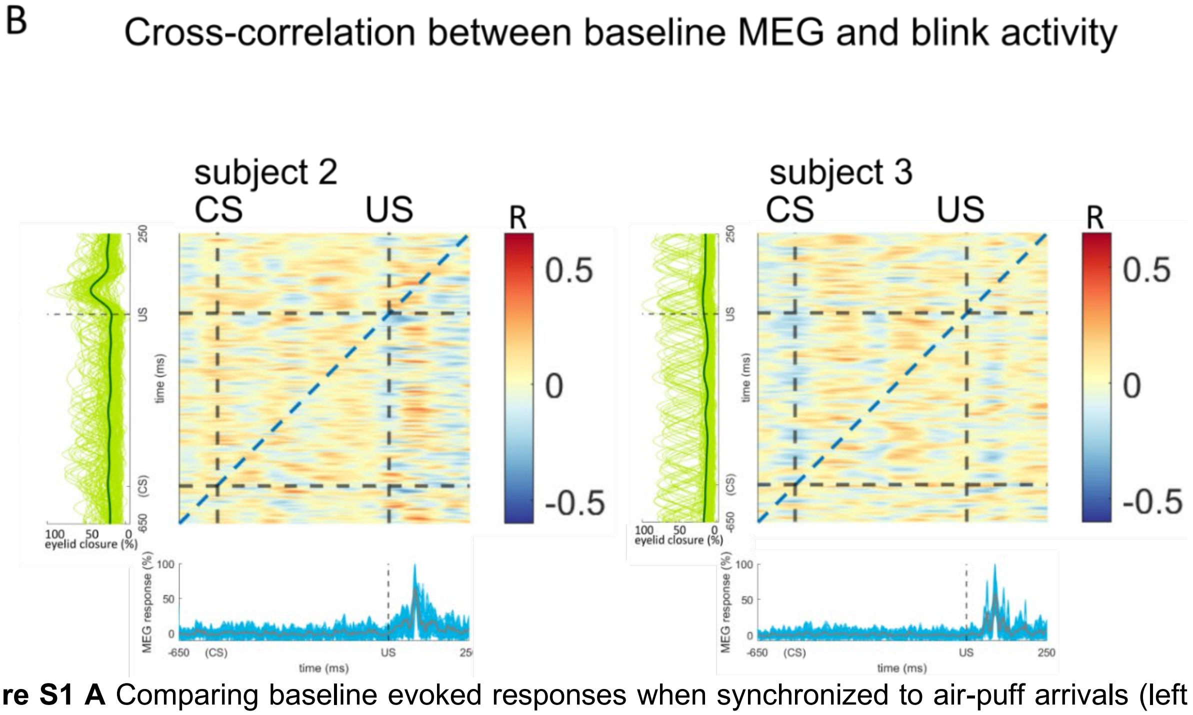


**Figure S1 A** Comparing baseline evoked responses when synchronized to air-puff arrivals (left column) or blink onsets (right). Response amplitude diminished after being re-aligned to blink onset, suggesting that they are better driven by the US. **B** Correlation matrices showed between-trial MEG (x-axis) and blink (y-axis) data interdependence at each timepoint between -650~250 ms, baseline phase. We used rectified data from each subject-specific maximum MEG channel to compute correlation matrices. Single-trial MEG (cyan curve) and average MEG (brown curve) waveform are presented horizontally, below the correlation matrix. Single-trial (light green) and average (dark green) blink trace are presented left to the correlation matrix of each participant. Here data from participants 2 &3 are shown, the correlation matrix of participants 1 & 4 are similar to presented data. Strong blink related responses would show up red areas along the diagonal line, which is not observed in our participants. The dashed vertical and horizontal lines indicate the CS-onset (-533 ms) and US-onset (0 ms).

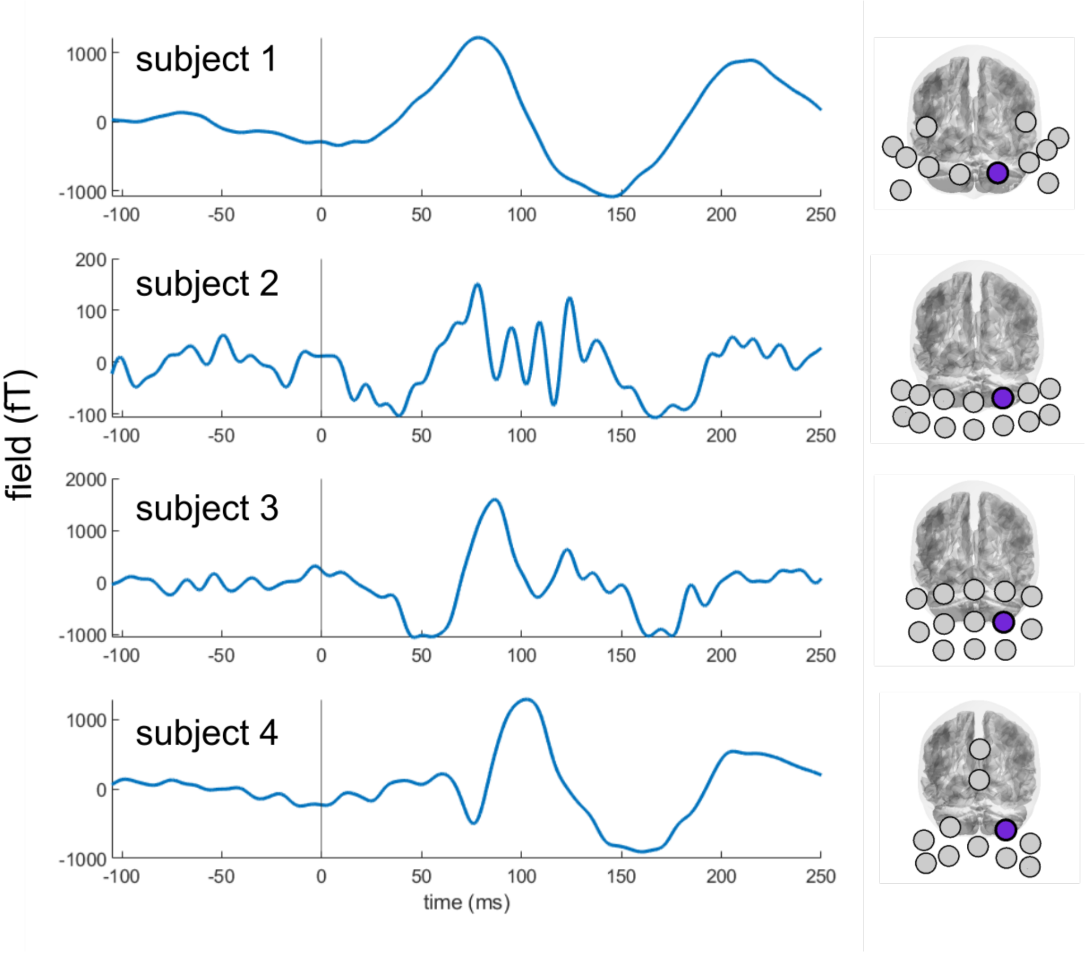


**Figure S2** Average waveforms of baseline (US only) MEG data from one channel of each participant. Channels were selected based on their sensor positions, to be close to the predicted locus of cerebellar responses, as seen in the right panel in blue circles. The time axis is relative to US onset.

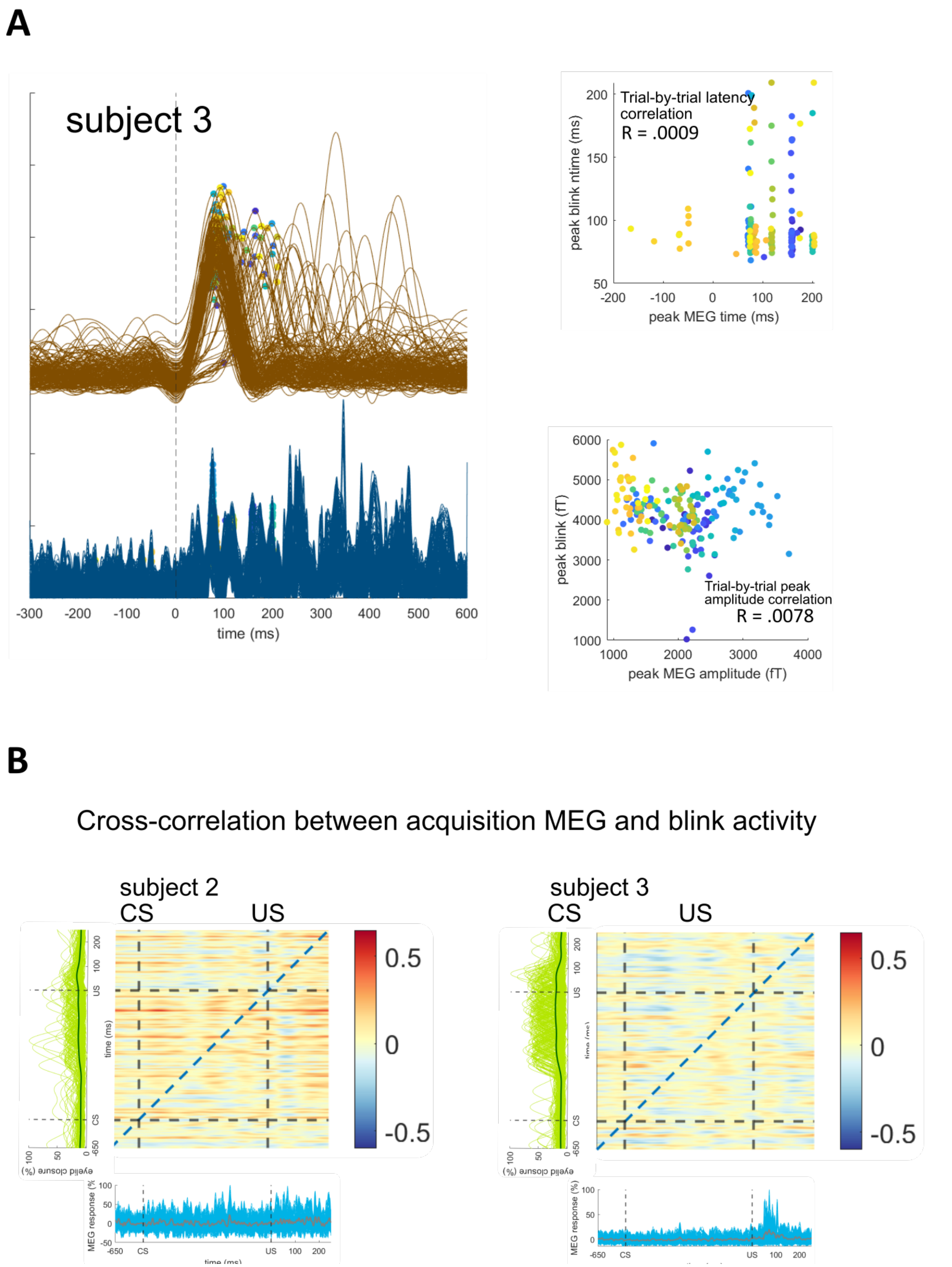


**Figure S3 A** Participant 3's CR(+) trial data of eyeblink (brown curves) and MEG (blue curves) in the acquisition phase. There was no trial-by-trial correlation of peak latency (right upper panel) nor of peak amplitude (right lower panel). **B** Correlation matrices show between-trial MEG (x-axis) and blink (y-axis) data interdependence at each timepoint between -650~250 ms, acquisition CR(+) trials. Participants 2 & 3's data are presented here. Rectified data from each subject-specific maximum MEG channel was used to compute correlation matrices. Strong correlation between MEG and blink responses would present as red areas above the diagonal line, which is not observed in our participants.

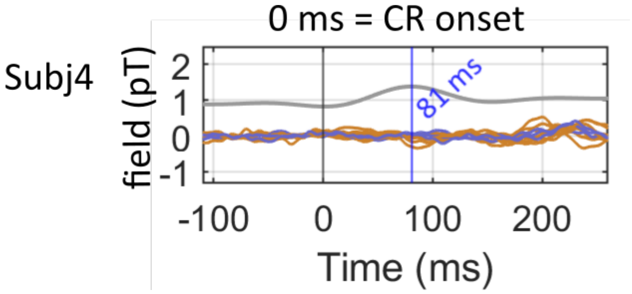


**Figure S4** Participant 4's average MEG and blink responses of CR+ trials in the acquisition phase. Trial data were aligned to the onset of CRs as for the other three participants (**Figure 4E, main text)**. Each colour trace corresponds to the average signal for one sensor over the posterior cranium, situated left (blue curves) and right (orange curves) of the midline respectively. Average blink peaked at 81 ms (blue text). No MEG peak was found around blink peak latency, different from what was seen in the data of three other participants (**Figure 4E**).
